# Identification of the alternative sigma factor regulons of *Chlamydia trachomatis* using multiplexed CRISPR interference

**DOI:** 10.1101/2023.04.27.538638

**Authors:** Nathan D. Hatch, Scot P. Ouellette

## Abstract

*C. trachomatis* is a developmentally regulated, obligate intracellular bacterium that encodes three sigma factors: σ66, σ54, and σ28. σ66 is the major sigma factor controlling most transcription initiation during early and mid-cycle development as the infectious EB transitions to the non-infectious RB that replicates within an inclusion inside the cell. The roles of the minor sigma factors, σ54 and σ28, have not been well characterized to date – however, there are data to suggest each functions in late-stage development and secondary differentiation as RBs transition to EBs. As the process of secondary differentiation itself is poorly characterized, clarifying the function of these alternative sigma factors by identifying the genes regulated by them will further our understanding of chlamydial differentiation. We hypothesize that σ54 and σ28 have non-redundant and essential functions for initiating late gene transcription thus mediating secondary differentiation in *Chlamydia*. Here, we demonstrate the necessity of each minor sigma factor in successfully completing the developmental cycle. We have implemented and validated multiplexed CRISPRi techniques novel to the chlamydial field to examine effects of knocking down each alternative sigma factor individually and simultaneously. In parallel, we also overexpressed each sigma factor. Altering transcript levels for either or both alternative sigma factors resulted in a severe defect in EB production as compared to controls. Furthermore, RNA sequencing identified differentially expressed genes during alternative sigma factor dysregulation, indicating the putative regulons of each. These data demonstrate the levels of alternative sigma factors must be carefully regulated to facilitate chlamydial growth and differentiation.

**Importance:** *Chlamydia trachomatis* is a significant human pathogen in both developed and developing nations. Due to the organism’s unique developmental cycle and intracellular niche, basic research has been slow and arduous. However, recent advances in chlamydial genetics have allowed the field to make significant progress in experimentally interrogating the basic physiology of *Chlamydia*. Broadly speaking, the driving factors of chlamydial development are poorly understood, particularly regarding how the later stages of development are regulated. Here, we employ a novel genetic tool for use in *Chlamydia* while investigating the effects of dysregulating the two alternative sigma factors in the organism that help control transcription initiation. We provide further evidence for both sigma factors’ essential roles in late-stage development and their potential regulons, laying the foundation for deeper experimentation to uncover the molecular pathways involved in chlamydial differentiation.

## Introduction

*Chlamydia trachomatis* (Ctr), like all *Chlamydiae*, is an obligate intracellular bacterium that utilizes a biphasic developmental cycle to alternate between infectious elementary bodies (EBs) and replicating reticulate bodies (RBs). During the infection cycle, an EB binds to a host cell and secretes effector proteins through a type-III secretion system (T3SS), inducing endocytosis of the EB by the host cell. Upon entry, the EB begins primary differentiation into an RB while secreting additional effectors that insert into the membrane of the newly formed endosome. These additional effectors serve diverse functions, most notably to divert the Ctr-containing endosome from the endolysosomal pathway and to promote its trafficking to a perinuclear location. This modified endosome is termed an inclusion – where Ctr will reside until the end of the developmental cycle. Altogether, these initial events are known as the early cycle. Mid cycle begins following primary differentiation as the RB rapidly multiplies, filling the inclusion with hundreds more RBs. Finally, late cycle is characterized by asynchronous secondary differentiation of RBs into EBs, followed by lysis of the inclusion and host cell, releasing the EBs to infect new host cells (reviewed extensively in (1, 2)). To accommodate the functional requirements of each stage, Ctr upregulates subsets of genes in a temporally regulated manner (3, 4). Importantly, the mechanisms governing this temporal regulation have not been fully delineated.

Sigma factors are required to guide prokaryotic RNA polymerase (RNAP) to a gene’s promoter before initiating transcription. The promoter sequence recognized is dependent on the sigma factor bound to RNAP. For example, σ70 of *E. coli* recognizes a different promoter sequence than σ32, and thus regulates a different subset of genes (5–7). Importantly, sigma factors are subdivided into two evolutionarily distinct families: the σ70 family, whose members exhibit high structural homology to canonically described housekeeping sigma factors, and the σ54 family, whose members require an ATP-dependent enhancer binding protein (EBP) to fully initiate transcription (8). Alternative sigma factors within the σ70 family are often regulated by anti-sigma factors, proteins that sequester or release an alternative sigma factor under certain conditions. This is in stark contrast to the σ54 family, members of which rely on an EBP to catalyze the formation of the transcription open complex. In general, EBPs are part of a two-component system including a sensor kinase and a response regulator, the EBP. The signals recognized by these systems vary, but often relate to stress response, sporulation, or other forms of developmental regulation (9). Furthermore, alternative sigma factors actively compete for access to RNAP. Changes in the overall availability of sigma factors through partner switching mechanisms, regulatory proteolysis, or gene expression, can skew the relative abundance of specific sigma factors, allowing dynamic competition between sigma factors to compete for RNAP. This is further influenced by the affinity and intrinsic dissociation constant of a given alternative sigma factor, meaning direct stoichiometric comparisons are not reliable in determining the competitiveness of a given alternative sigma factor (10).

As previously noted, chlamydial gene expression is broadly characterized into three temporal categories: early, mid, and late cycle (3, 4). The general hypothesis that temporal gene regulation is driven by alternative sigma factors has been posited for decades (11, 12); however, the mechanism by which this occurs, if at all, is unknown. Ctr encodes three sigma factors: i) σ66 of the σ70 family, the primary sigma factor responsible for most transcription initiation in Ctr, ii) σ28 of the σ70 family, hypothesized to regulate a subset of late genes (13), and iii) σ54 of the σ54 family, also hypothesized to regulate a separate subset of late genes, particularly those related to the T3SS (14). Furthermore, Ctr encodes a two-component system, AtoC/S, where AtoC serves as an EBP and AtoS as its sensor kinase (15, 16). This chlamydial two-component system has also been referred to as NtrB/C and CtcB/C. While multiple studies have utilized *in silico* and *in vitro* models (either heterologous or organism-free) to suggest the σ28 and σ54 regulons in Ctr, only one recent study has provided *in vivo* (defined herein as “in the chlamydial organism”) evidence (12–14). Given our curiosity of each sigma factor’s respective roles in transcription, we sought to complement existing evidence with a broader *in vivo* approach utilizing RNA sequencing (RNA-seq) in combination with either inducible CRISPR interference (CRISPRi) mediated knockdown or inducible overexpression to further elucidate these alternative sigma factor regulons and their developmental role in Ctr. Furthermore, to investigate whether Ctr’s alternative sigma factors exhibit any cooperative function, we adapted a multiplex CRISPRi technique for simultaneous knockdown of multiple gene targets for use within Ctr. The DNase-dead *Acidoaminococcus* dCas12 ortholog, ddCpf1, has been successfully used in multiplexed knockdown studies outside of *Chlamydia* (17) and has been recently adapted for use in Ctr – although only for single target knockdown (18). By understanding the extent to which alternative sigma factors control transcription, we can more easily direct future studies. This direction is critical to pursue the molecular mechanisms governing fundamental processes such as temporal gene regulation and differentiation. Here, we demonstrate the first use of multiplexed CRISPRi knockdown in Ctr and provide further *in vivo* evidence of each alternative sigma factor’s regulons.

## Results

### Development of inducible overexpression and knockdown strains in Ctr

To expand upon the existing knowledge based primarily on *in silico* and *in vitro* studies and to characterize the regulons of sigma factors 28 and 54 in *Chlamydia*, we developed a variety of inducible plasmid constructs for use *in vivo* (Table 1). To limit leaky expression, all plasmids were developed using the pBOMBL backbone as described previously (18). We generated ectopic overexpression constructs encoding σ28_10xHistidine (*fliA*), σ54_10xH (*rpoN*), σ66_10xH (*rpoD*), and a dual overexpression vector encoding polycistronic σ28_FLAG and σ54_10xH. Additionally, we developed knockdown constructs by inserting various crRNAs into the pBOMBL12CRia (pL12CRia) vector described previously (18). These included a non-targeting crRNA (i.e., a negative control with no Ctr homology) or crRNAs targeting the promoter regions of σ28, σ54, or *incA* either individually or as a multiplexed combination for two or three targets. The triple knockdown construct was created as a proof-of-concept utilizing a previously characterized *incA* crRNA known to be effective when used individually (18). The resulting plasmids were transformed into Ctr serovar L2.

**Table 1.** Plasmids constructed and used in this study. The pBOMBL backbone is used for overexpression, while pL12CRia is used for knockdown. Both backbones produce GFP and beta-lactamase.

| Constructs Utilized |
| --- |
| pBOMBL-$\sigma_{28}$ 10xHistidine |
| pBOMBL-$\sigma_{54}$ 10xHistidine |
| pBOMBL-$\sigma_{28}$ FLAG + $\sigma_{54}$ 10xHistidine |
| pBOMBL-$\sigma_{66}$ 10xHistidine |
| pL12CRia (NT) |
| pL12CRia (σ28) |
| pL12CRia (σ54) |
| pL12CRia (σ28+σ54) |
| pL12CRia (σ28+σ54+incA) |

To validate these new strains, RT-qPCR was performed to analyze specific transcriptional changes associated with overexpression or knockdown of selected genes. Briefly, McCoy cells were infected with each strain and induced or not with 10 nM anhydrotetracycline (aTc). Knockdown strains were induced at 4 hours post infection (hpi) and collected at 24 hpi. Induction at 4 hpi ensures sustained knockdown prior to initial sigma factor transcription, which occurs as early as 8 hpi (3, 11). Overexpression constructs were induced at 14 hpi and collected at 18 hpi. This window was selected with RNA-seq in mind, seeking to maximize contrast between uninduced and overexpression strains before potential secondary effects accumulate. Upon induction, transcripts were altered as predicted (Figure 1). For example, *σ28* transcripts were significantly reduced ~75-80% when *σ28* was targeted for knockdown (either individually or multiplexed) whereas those transcripts were increased approximately 30-fold when σ28 was overexpressed (either individually or in combination with σ54). Surprisingly, *σ66* transcripts were upregulated 30-fold on average when σ54_10xH was overexpressed while the opposite was observed during induction of σ28_FLAG and σ54_10xH together, as *σ66* transcripts were reduced 2-fold compared to uninduced (Figure 1B). The reason for the upregulation of *σ66* transcripts during σ54 overexpression alone is not apparent. Most notably, knockdown efficiencies for each sigma factor measured in the strains carrying the multiplexed pL12CRia(σ54/σ28)::L2 or pL12CRia(incA/σ54/σ28)::L2 plasmids were similar to knockdown in strains targeting the individual gene (e.g., pL12CRia(σ28)), demonstrating the success of multiplexed CRISPRi in Ctr. Importantly, this indicates that the addition of targets does not compromise knockdown efficiency since the triple knockdown strain showed the same level of repression for the sigma factors as the dual or single knockdown strain (Figure 1A). These data validate that our overexpression and knockdown strains exhibited the expected transcript changes associated with the targeted sigma factor.

**Figure 1.**
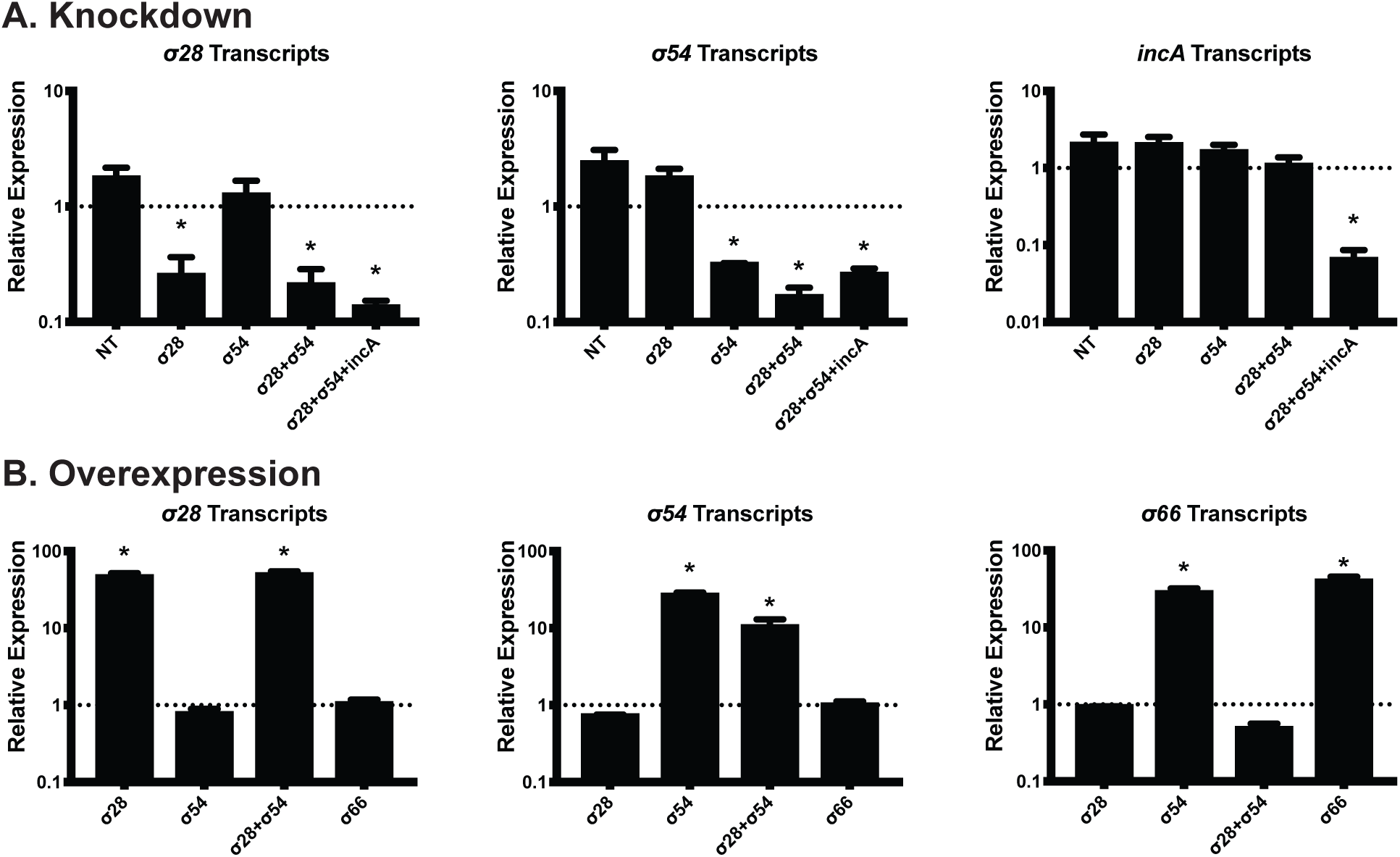
Validation of knockdown and overexpression strains via RT-qPCR. A) Knockdown strains (x-axis) were induced or not with 10 nM aTc at 4 hpi. Samples were collected at 24 hpi and processed as described in the methods. Uninduced ratios of cDNA:16S are set to 1, induced samples’ cDNA:16S ratios are shown as a relative proportion. NT: Non-Targeting. B) Overexpression strains (x-axis) were induced or not with 10 nM aTc at 14 hpi and collected at 18 hpi. All samples include three biological replicates. * = p<0.05 via ratio paired t-test.

### Dysregulation of alternative sigma factors 28 and 54 is detrimental to developmental cycle progression

Given the temporal nature of chlamydial gene regulation, we reasoned that any deviation in sigma factor activity would be detrimental to chlamydial development. To assess this hypothesis, we performed inclusion forming unit (IFU-a proxy for infectious EBs) assays to determine each strain’s ability to produce infectious progeny and immunofluorescence assays (IFA) to examine organism morphology using the same induction conditions as described above. As the triple knockdown strain carrying the pL12CRia(incA/σ54/σ28)::L2 plasmid was created as a proof-of-principle, we did not include it for further analysis. Knockdown of either or both alternative sigma factors resulted in a greater than 2-log decrease in infectious progeny at 24 hpi (Figure 2A). Similarly, overexpressing σ28 with or without simultaneously overexpressing σ54 resulted in a greater than 1.5-log decrease in infectious progeny (Figure 2B). Interestingly, σ54 overexpression alone produced an ~80% decrease in IFUs. Considering our model does not include any manipulation of the AtoS/AtoC two-component system required for σ54 to initiate transcription, we speculate that this decrease in IFUs is attributable to the increase in *σ66* transcripts (Figure 1B), as σ66 overexpression alone results in a ~70% decrease in IFUs.

**Figure 2.**
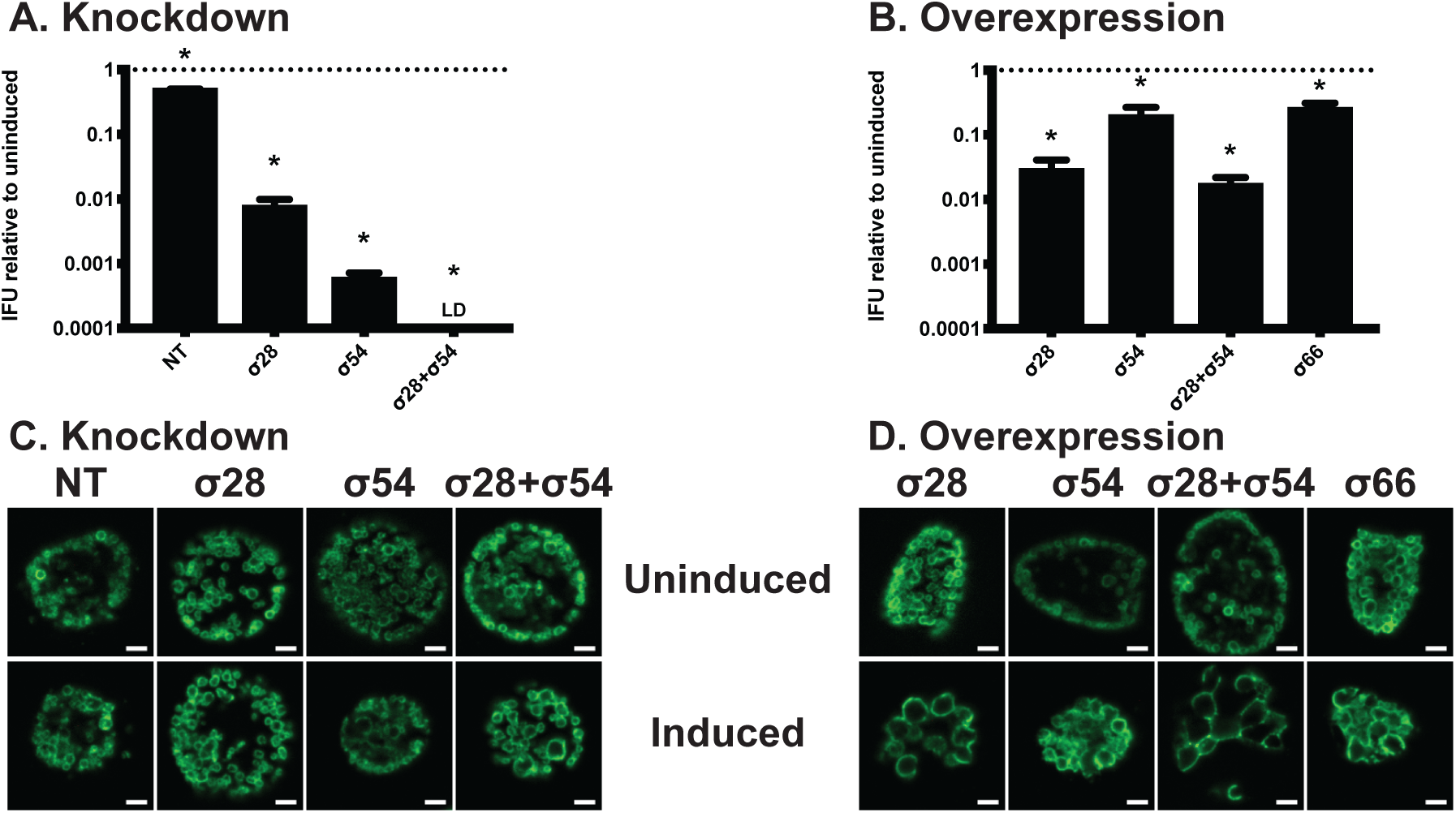
Sigma factor dysregulation is detrimental to developmental cycle completion. Inclusion Forming Unit (IFU) assays were performed to assess the impact of sigma factor knockdown (A) or overexpression (B) on the ability to produce infectious progeny. Strains were induced or not with 10 nM aTc at 4 hpi (A) or 14 hpi (B) and collected at 24 hpi for reinfection and enumeration. Values are expressed as a ratio of EBs extracted from induced samples and their respective uninduced counterparts. NT: Non-Targeting. All samples include three biological replicates. * = p<0.05 via ratio paired t-test. (C-D) Immunofluorescence assay (IFA) was performed to assess inclusion size and organism morphology using the same induction conditions described in A-B. At 24 hpi, cells were fixed with methanol and stained with primary antibodies to Ctr’s Major Outer Membrane Protein (MOMP). Representative images of three biological replicates are shown. All images were acquired on an AXIO Imager.Z2 with ApoTome.2 at 100x magnification. Scale bars represent 2 µm.

Under normal conditions, chlamydial RBs are ~1 µm in diameter while EBs are ~0.3 µm. RB size can be significantly larger under various conditions of stress, such as nutrient deprivation, antibiotic exposure, etc. ((19–24)). Interestingly, the knockdown strains appeared morphologically normal with no obvious change compared to the non-targeting strain (Figure 2C). The exception was knockdown of both σ28 and σ54 that showed some slightly enlarged RB forms. Overall, the relatively normal morphology of the knockdown strains stood in contrast to their significant drop in infectivity (Fig. 2A). Unsurprisingly, we qualitatively observed the most notable morphological aberrations with the overexpression strains – namely, those overexpressing σ28 (Figure 2D). These data demonstrate the dispensability of the alternative sigma factors for early and mid-cycle development (given the normal morphology when knocked down) and their indispensability in late-cycle development (given the reduction in IFUs), indicating their requirement for successful secondary differentiation.

### Analysis of RNA-seq data from overexpression and knockdown strains

To obtain unbiased transcriptome data, RNA-seq was performed on three biological replicates using induced and uninduced culture conditions from each strain with RNA samples collected at 24 hpi (knockdown strains) or 18 hpi (overexpression strains). Prior to sequencing, RNA was processed to enrich bacterial mRNA, as described previously (25), using a combination of Oligotex, MicrobEnrich, and MicrobExpress. Samples were submitted to the UNMC Genomics core facility for further processing, library generation, and sequencing. On average, each sample had approximately 600k reads map to Ctr L2. After accounting for rRNA contamination and read length, we estimate ~30-fold genome coverage, suggesting adequate read depth per sample (Table 2). Statistical analyses between uninduced and induced samples were performed by the UNMC Bioinformatics Core Facility. Significant results were defined as having 2-fold change, p-value < 0.05, and false discovery rate (FDR) < 0.05.

**Table 2.** Generalized representation of sequencing efficiency. While the percentage of reads mapped to Ctr is low (<2%), the raw number of reads mapped to Ctr provide ample coverage (est. >30-fold). Estimated genome coverage calculated by multiplying reads mapped to *Chlamydia* by 100 (bp per read), multiplying by % Ctr reads - protein coding, and dividing by bp in the genome (1,038,842).

|  | Average per sample |
| --- | --- |
| Number of reads | 36,400,449 |
| Reads mapped to Ctr | 609,745 |
| % of reads mapped to Ctr | 1.68 |
| % Ctr reads - protein coding | 98.40 |
| % Ctr reads - rRNA | 1.59 |
| Est. genome coverage (fold) | 33.06 |

Additionally, for knockdown samples, a second analysis was performed to determine statistical significance between the induced non-targeting strain and each induced knockdown strain. The purpose of this was to account for any changes in transcription due to the expression of dCas12 independent from targeted knockdown. We previously observed that dCas12 expression in combination with the NT crRNA can slightly delay developmental cycle progression, resulting in ~2-fold lower IFU yields while having no impact on genomic DNA levels or bacterial morphology (26). For knockdown samples, significant results were cross-referenced between “induced vs uninduced” and “induced vs induced non-targeting” and only results significant in both sets were analyzed further. Although the non-targeting crRNA sequence showed no homology to any Ctr L2 sequence, the RNA-seq analysis indicated that *lepA* transcripts were significantly reduced after inducing dCas12 expression, but only in the non-targeting strain. This was surprising given that there is no homology to the *lepA* coding or intergenic sequence and no PAM sequence that would direct a low-homology overlap to the gene. When measured by RT-qPCR, *lepA* transcripts were reduced ~3-fold during induction of dCas12 expression, but only in the strain co-expressing the NT crRNA (Suppl. Figure 1). Notably, *lepA* transcripts were not decreased in the empty vector (dCas12 with no crRNA). For future studies, we recommend that the empty vector be utilized as a negative control instead of a non-targeting strain to limit random effects associated with expressing an irrelevant crRNA sequence.

### The σ28 regulon is composed of two genes

As indicated above, our goal was to define the regulons of the alternative sigma factors. To date, there have been no *in vivo* studies to experimentally determine the σ28 regulon. Surprisingly, knockdown of *σ28* resulted in significant loss of only *hctB* and *tsp* transcripts at a magnitude of 8 and 16-fold, respectively (Table 3). Both genes have been predicted to be σ28-regulated in *in vitro* studies (13, 27). Furthermore, both genes were upregulated in the *σ28_10xH* overexpression strain (21 and 191-fold, respectively), strengthening the case that these two genes are the only genes regulated by σ28. Nonetheless, in addition to *hctB* and *tsp*, 13 other genes were upregulated between 2 and 4-fold following σ28_10xH expression only and not following knockdown of *σ28* (Suppl. Table 1). We cross-referenced the upregulated genes with σ28’s predicted regulon (published by Yu et al. (13)) and found 3 of the 9 predicted genes were upregulated in our overexpression sample – *hctB, tsp,* and *yebL* (*ct415/ctl0672*). Notably, while *yebL* was found to be significantly upregulated ~2-fold by RNA-seq, *hctB* and *tsp* were upregulated 21 and 191-fold, respectively. The disparity between *hctB/tsp* and *yebL* suggests the latter is not a specific target of σ28. Rather, the changes in transcript levels during overexpression may reflect differences in the developmental cycle (i.e., an indirect effect of overexpression). Overall, these data indicate that σ28 specifically regulates only *hctB* and *tsp* – an unexpected finding.

**Table 3.** All significant hits observed during σ28 knockdown (KD) are listed alongside the fold changes observed during overexpression (OE). “Name” includes the annotated name under the accession file used for annotation. Other common names may be given in ( ). Y: Yu et al. found these to be regulated by σ28 *in vitro*.

| CT# | CTL# | Name | Function | Fold change |  |  |
| --- | --- | --- | --- | --- | --- | --- |
| | | | | σ28 KD | σ28 OE | |
| CT441 | CTL0700 | tsp | Tail-specific protease | -15.39 | 191.24 | Y |
| CT061 | CTL0317 | fliA (σ28) | Sigma factor | -13.23 | 78.58 | |
| CT046 | CTL0302 | hct2 (hctB) | Histone-like protein 2 | -7.90 | 21.48 | Y |

### σ54 is epistatic to σ28

Recent *in vivo* experimentation carried out by Soules et al. utilized a σ54 hyperactivation model to investigate σ54-dependent transcriptional changes in Ctr (14). This provided a unique opportunity to implement our knockdown techniques to validate and complement their study. Knockdown of σ54 had a much broader impact on transcription than σ28 knockdown, significantly downregulating 67 genes in our dataset. Downregulated genes were cross-referenced with genes identified by both Soules et al. and Mathews and Timms (12, 14). Interestingly, none of the nine genes predicted by Mathews and Timms appeared in our results. We were, however, able to recapitulate approximately half of the predictions made by Soules et al. Notably, five of our unique hits are generically classified as secretion-associated, matching the broad regulatory theme proposed by Soules et al. Beyond secretion-associated genes, unique hits of interest also include transcription factors *dksA* and *chxR.* Unfortunately, DksA’s role in chlamydial gene regulation is still largely unknown and therefore cannot be directly linked to any potential genes in the σ54 regulon. Conversely, three genes from ChxR’s five-gene regulon also appear in our data: *ctl0466* (*ct214*), *ctl0828* (*ct565*), and *ctl0063* (*ct694*). This is unsurprising, considering ChxR is a transcription activator, and reduction in ChxR would logically result in a reduction of its associated regulon. Furthermore, despite *chxR* not being present in the Soules et al. data set, two of the three ChxR regulated genes detected in our data set were detected in theirs: *ctl0828* and *ctl0063*. Most notably, *tsp* and *hctB* appear to be downregulated upon σ54 knockdown. While *tsp* was a unique hit to our study, Soules et al. also detected *hctB.* This suggests σ54 activity is a necessary precursor for endogenous σ28 activity.

In contrast, our σ54 overexpression samples did not demonstrate broad transcriptional changes. Considering σ54 requires an active EBP to fully initiate transcription, which our model does not include, it is likely that overexpression of σ54 does not correlate with an increase in transcriptional activation due to the lack of a concurrently active EBP. Nonetheless, two genes were significantly altered: *σ66* and *euo* (Suppl. table 2). Given our qPCR analysis (Fig. 1), we expected to find *σ66* upregulated in this sample. We speculate that the increase in *euo* transcripts may be due to excessive σ66 production and an associated delay in developmental progression. While σ66 overexpression had a noticeable effect on infectious progeny production, there was only 1 significant transcriptional change: a 2-fold increase in *euo.* This is to be expected, as the natural amount of σ66 is expected to outnumber RNAP 3:1 and thus not pose as a rate-limiting factor of transcription (28). Therefore, increasing σ66 amounts would not be expected to cause an increase in σ66-dependent transcription. However, the mechanism for excess σ66 reducing IFUs by 80% without inducing significant transcriptional changes beyond *euo* and *σ66* is not clear. Due to the differing experimental endpoints, it is possible that sequencing at 24 hpi would reveal significant deficiencies in late genes, as excess σ66 could be outcompeting σ28 and σ54 – this would explain the decrease in IFUs seen at 24 hpi.

### Multiplex knockdown reveals an expansion of significant hits compared to single knockdown

Development of a dual knockdown technique for both alternative sigma factors was pursued in order to assess the potential for cooperative effects. Analysis of our σ28 + σ54 double knockdown revealed an intriguing expansion of differentially expressed genes in addition to those identified during single knockdown. Importantly, 62 of 67 significantly downregulated genes from σ54 knockdown were recapitulated in the double knockdown, including the proposed σ28 regulon of *tsp* and *hctB*. Furthermore, 27 additional genes were significantly downregulated upon double knockdown, including several outer membrane proteins and T3SS proteins, conforming to the trend observed by Soules et al. (14). Broadly, we identified 18 T3SS associated genes, 10 associated with gene regulation, 7 outer membrane components, and 9 metabolism related genes (Tables 4–8). Those categories may be underestimates, as there are 41 uncharacterized hypothetical genes that could not be assigned a functional category (Suppl. table 3). Particularly noteworthy are the decreases in specific T3SS structural genes such as *fliI* and *flhA,* two potentially redundant genes that are homologous to genes found in flagellating bacteria. Of the “gene regulation” subset, *rsbU* and *rsbV1* are downregulated, suggesting σ66 activity may be reduced downstream (29). Furthermore, *mcsA/B* regulate gene expression at the protein level through modulating degradation via the ClpC system – this may be a significant driver of the proteomic shift that occurs during secondary differentiation (30). Additionally, *ispA/F* and *sucB1/C* are significantly downregulated, suggesting isoprenoid biosynthesis and succinate metabolism, respectively, are important for late-stage development and influenced by σ54 activity.

**Table 4.** T3SS-associated genes detected during dual knockdown (KD) of σ28 + σ54 and the accompanying fold changes observed. Also included are fold changes during σ54 knockdown alone and σ28 + σ54 dual overexpression (OE) for those genes. “Name” includes the annotated name under the accession file used for annotation. Other common names may be given in ( ). S: Soules et al. observed upregulation of these genes in a constitutively active AtoC model, mimicking σ54 hyperactivity. W: These genes are upregulated during σ28 + σ54 OE but are not observed in σ28 OE.

| CT# | CTL# | Name | Function | Fold change |  |  |  |
| --- | --- | --- | --- | --- | --- | --- | --- |
| | | | | $\sigma 28/\sigma 54$<br>KD | $\sigma 54$<br>KD | $\sigma 28/\sigma 54$<br>OE | |
| CT576 | CTL0839 | scc2 (lcrH 1) | T3SS chaperone | -16.94 | -6.88 | - | S |
| CT579 | CTL0842 | copD | T3SS translocator | -15.54 | -6.15 | - | S |
| CT578 | CTL0841 | copB | T3SS translocator | -12.07 | -5.69 | - | S |
| CT456 | CTL0716 | tarp | Translocated actin-recruiting phosphoprotein | -7.18 | -3.93 | - | S |
| CT717 | CTL0086 | fliI | T3SS component, ATPase | -3.76 | - | - |  |
| CT005 | CTL0260 | CTL0260 (incV) | Inclusion membrane protein | -3.35 | - | - | S |
| CT719 | CTL0088 | fliF | T3SS component, M-ring | -3.35 | -3.09 | - |  |
| CT712 | CTL0081 | CTL0081 | T3SS effector | -3.35 | -2.26 | 2.27 | W |
| CT694 | CTL0063 | CTL0063 | T3SS effector | -3.19 | -2.85 | 2.32 | S/W |
| CT214 | CTL0466 | CTL0466 | Inclusion membrane protein | -3.03 | -2.70 | - |  |
| CT564 | CTL0827 | sctT | T3SS component, export apparatus | -2.92 | -2.13 | - |  |
| CT562 | CTL0825 | sctR | T3SS component, export apparatus | -2.60 | -2.52 | - |  |
| CT060 | CTL0316 | flhA | T3SS component, export gate | -2.52 | - | - |  |
| CT274 | CTL0526 | CTL0526 | T3SS chaperone | -2.31 | - | - |  |
| CT090 | CTL0345 | lcrD | Low calcium response protein D | -2.28 | - | - |  |
| CT561 | CTL0824 | sctL | T3SS component, spoke protein | -2.28 | - | - |  |
| CT559 | CTL0822 | sctJ | T3SS component, inner ring | -2.24 | -2.01 | - |  |
| CT089 | CTL0344 | lcrE (copN) | Low calcium response protein E | -2.24 | - | - |  |

**Table 5.** Genes associated with gene regulation, either directly or through regulating proteolysis pathways, detected during dual knockdown (KD) of σ28 + σ54 and the accompanying fold changes observed. Also included are fold changes during σ54 knockdown alone and σ28 + σ54 dual overexpression (OE) for those genes. “Name” includes the annotated name under the accession file used for annotation. Other common names may be given in ( ).

| CT# | CTL# | Name | Function | Fold change |  |  |
| --- | --- | --- | --- | --- | --- | --- |
| | | | | $\sigma 28/\sigma 54$<br>KD | $\sigma 54$<br>KD | $\sigma 28/\sigma 54$<br>OE |
| CT609 | CTL0873 | rpoN ($\sigma 54$) | Sigma factor | -26.87 | -21.81 | 35.37 |
| CT061 | CTL0317 | fliA ($\sigma 28$) | Sigma factor | -9.70 | - | 82.07 |
| CT424 | CTL0683 | rsbV | Anti-sigma F factor antagonist | -5.03 | -4.26 | - |
| CT589 | CTL0852 | CTL0852 | PP2C-like phosphatase | -4.83 | -4.17 | - |
| CT675 | CTL0044 | aspC_2<br>(mcsB) | Protein-arginine kinase | -3.93 | - | - |
| CT630 | CTL0894 | CTL0894<br>(chxR) | Transcription regulator | -3.56 | -2.20 | - |
| CT467 | CTL0727 | atoS (ctcB) | Sensor histidine kinase | -3.28 | -2.38 | - |
| CT676 | CTL0045 | CTL0045<br>(mcsA) | Protein-arginine kinase<br>activator protein | -2.71 | -2.27 | - |
| CT407 | CTL0664 | dksA | RNAP-binding transcription<br>factor | -2.60 | -2.48 | - |
| CT477 | CTL0738 | CTL0738 | DNA methyltransferase | -2.40 | - | - |
| CT588 | CTL0851 | rbsU (rsbU) | Serine phosphatase | -2.34 | - | - |
| CT335 | CTL0589 | CTL0589 | Nucleoid associated protein | -2.12 | - | - |

**Table 6.** Both genes presumed to be σ28 regulated appeared during dual knockdown (KD) of σ28 + σ54 and the accompanying fold changes observed. Also included are fold changes during σ54 knockdown alone and σ28 + σ54 dual overexpression (OE) for those genes. “Name” includes the annotated name under the accession file used for annotation. Other common names may be given in ( ). S: Soules et al. observed upregulation of these genes in a constitutively active AtoC model, mimicking σ54 hyperactivity.

| CT# | CTL# | Name | Function | Fold change |  |  |  |
| --- | --- | --- | --- | --- | --- | --- | --- |
| | | | | $\sigma 28/\sigma 54$<br>KD | $\sigma 54$<br>KD | $\sigma 28/\sigma 54$<br>OE | |
| CT441 | CTL0700 | tsp | Tail-specific protease | -15.04 | -7.55 | 833.45 |  |
| CT046 | CTL0302 | hct2 (hctB) | Histone-like protein 2 | -7.03 | -4.82 | 63.86 | S |

**Table 7.** Genes associated with metabolism detected during dual knockdown (KD) of σ28 + σ54 and the accompanying fold changes observed. Also included are fold changes during σ54 knockdown alone and σ28 + σ54 dual overexpression (OE) for those genes. “Name” includes the annotated name under the accession file used for annotation. Other common names may be given in ( ). S: Soules et al. observed upregulation of these genes in a constitutively active AtoC model, mimicking σ54 hyperactivity. W: These genes are upregulated during σ28 + σ54 OE but are not observed in σ28 OE.

| CT# | CTL# | Name | Function | Fold change |  |  |  |
| --- | --- | --- | --- | --- | --- | --- | --- |
| | | | | $\sigma 28/\sigma 54$<br>KD | $\sigma 54$<br>KD | $\sigma 28/\sigma 54$<br>OE | |
| CT133 | CTL0388 | ubiE | Demethylmenaquinone methyltransferase | -7.04 | - | - |  |
| CT374 | CTL0628 | CTL0628 | Arginine/agmatine antiporter | -4.51 | -3.37 | - |  |
| CT372 | CTL0626 | CTL0626 | Porin | -3.66 | -2.50 | - |  |
| CT434 | CTL0693 | ispF | Isoprenoid biosynthesis | -2.84 | -2.32 | - |  |
| CT489 | CTL0750 | glgC | Glucose-1-phosphate adenylyltransferase | -2.55 | -2.59 | - | S |
| CT055 | CTL0311 | sucB 1 | Succinyltransferase | -2.54 | - | - |  |
| CT417 | CTL0674 | CTL0674 | Metal ABC transporter permease | -2.35 | - | 2.02 | W |
| CT821 | CTL0193 | sucC | Succinyl-CoA ligase subunit beta | -2.27 | -2.10 | - |  |
| CT628 | CTL0892 | ispA | Farnesyl diphosphate synthase | -2.07 | - | - |  |

**Table 8.**
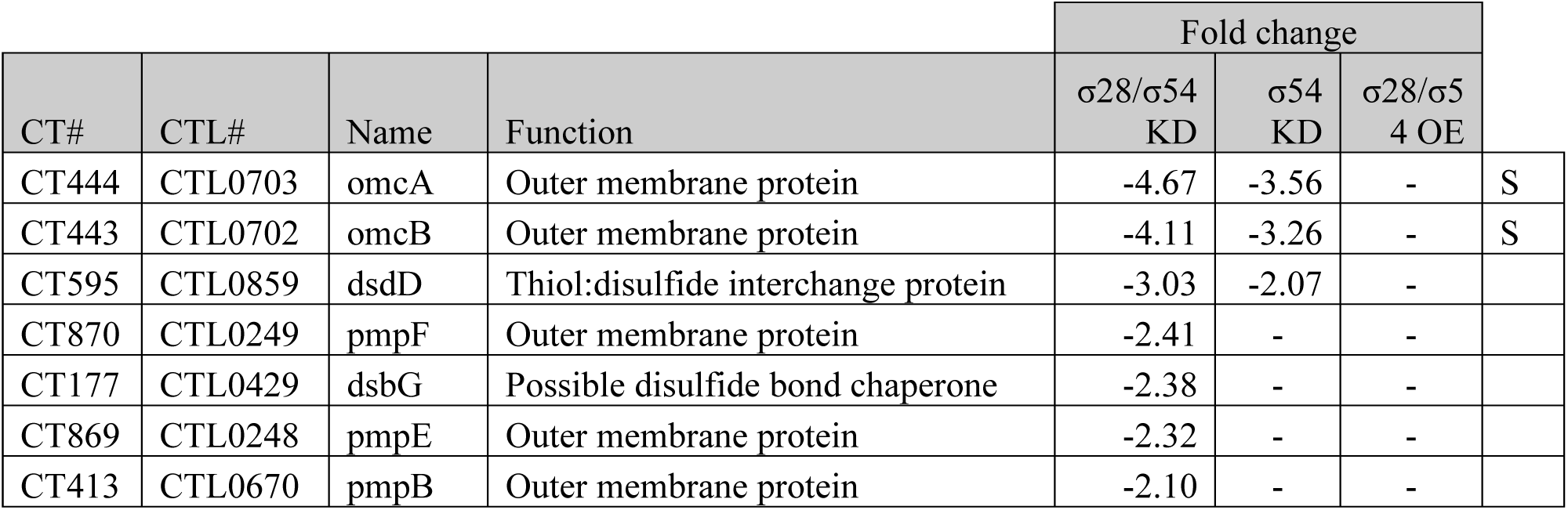
Genes associated with membrane organization detected during dual knockdown (KD) of σ28 + σ54 and the accompanying fold changes observed. Also included are fold changes during σ54 knockdown alone and σ28 + σ54 dual overexpression (OE) for those genes. “Name” includes the annotated name under the accession file used for annotation. Other common names may be given in ( ). S: Soules et al. observed upregulation of these genes in a constitutively active atoC model, mimicking σ54 hyperactivity.

Interestingly, dual overexpression of σ28 + σ54 resulted in an upregulation of 11 genes identified in the dual knockdown (9 from σ54 single knockdown). This is particularly surprising due to the lack of results observed from σ54 overexpression alone, save a 30-fold increase in *σ66* transcripts; notably, that increase does not occur during dual overexpression of σ28 + σ54. While 4 of the 11 genes of interest can be attributed to σ28 overexpression, the remaining 7 are suspected to be σ54 regulated (noted with W in Tables 4, 7, and supplemental table 3). Yet, as with the single σ54 overexpression system, additional σ54 produced during dual overexpression is not expected to be active. Nonetheless, these 7 genes appear to be indirectly regulated under circumstances of σ28 activity and increased σ54 levels, potentially through an indirect mechanism that involves decreasing σ66-bound RNAP. Importantly, this may be the same mechanism through which σ28 overexpression slightly upregulates genes not expected to be σ28-regulated.

## Discussion

*Chlamydia trachomatis* (Ctr) has long held the title of “most common bacterial sexually transmitted disease” for several reasons, one of them being the relative difficulty in producing scientific advancements in the field of chlamydial biology. Only within the last decade has the field developed the ability to transform Ctr – and more recently still the ability to transform other species within the genus (31–36). These milestone achievements have ushered in a “golden age” of chlamydial genetics, setting the foundation for adapting tools such as inducible expression, developmental reporter systems, allelic exchange, targeted knockdown, and more for use in Ctr (reviewed extensively in (32, 37)). Here, we expanded on this genetic toolbox by describing the first use of multiplexed knockdown mediated by CRISPRi in *Chlamydia*. Furthermore, we utilized this advancement to interrogate the effects of alternative sigma factor dysregulation in Ctr.

Our investigation provides further support for both σ28 and σ54 as late gene regulators essential for secondary differentiation. Knockdown of either is detrimental to the production of EBs while having no visible effect on chlamydial morphology, demonstrating their limited role in early and mid-cycle development. Moreover, genes affected by knockdown skew heavily towards late cycle genes. In agreement with a recent study published by Soules et al. (14), our σ54 knockdown detected several T3SS associated genes including structural components, chaperones, and effectors. Furthermore, we detected two key genes regulated by σ28 in the σ54 knockdown dataset, *hctB* and *tsp*, suggesting σ54 is epistatic to σ28 during the transition into late-cycle. In other organisms, σ28 regulates late-stage flagellar assembly; to prevent premature production of specific flagellar components, σ28 is sequestered by an anti-sigma factor (38–40). Following the completion of the hook and basal-body features of the flagella, a conformational change occurs to allow the anti-sigma factor to be secreted (38–40). The newly freed σ28 can then upregulate the remaining flagellar genes to complete the structure. Given the presence of several structural T3SS genes and the apparent dependency on σ54 for σ28 activity, we speculate that a similar mechanism exists in *Chlamydia* (Figure 3). It should be noted that our σ28 overexpression data do not conflict with this hypothesis, as overexpression would likely create a stoichiometric discrepancy between σ28 and the endogenous anti-sigma factor. While this hypothesis is highly speculative, meaningful progress could be made utilizing existing tools in combination with the direction provided by this study. Targeted knockdown and overexpression of single or multiple T3SS genes identified here followed by analysis of σ28-associated genes would be one potential avenue. While an anti-σ28 protein has yet to be identified, purification of chlamydial σ28 during mid-cycle development may reveal which protein acts as its anti-sigma factor, if any.

**Figure 3.**
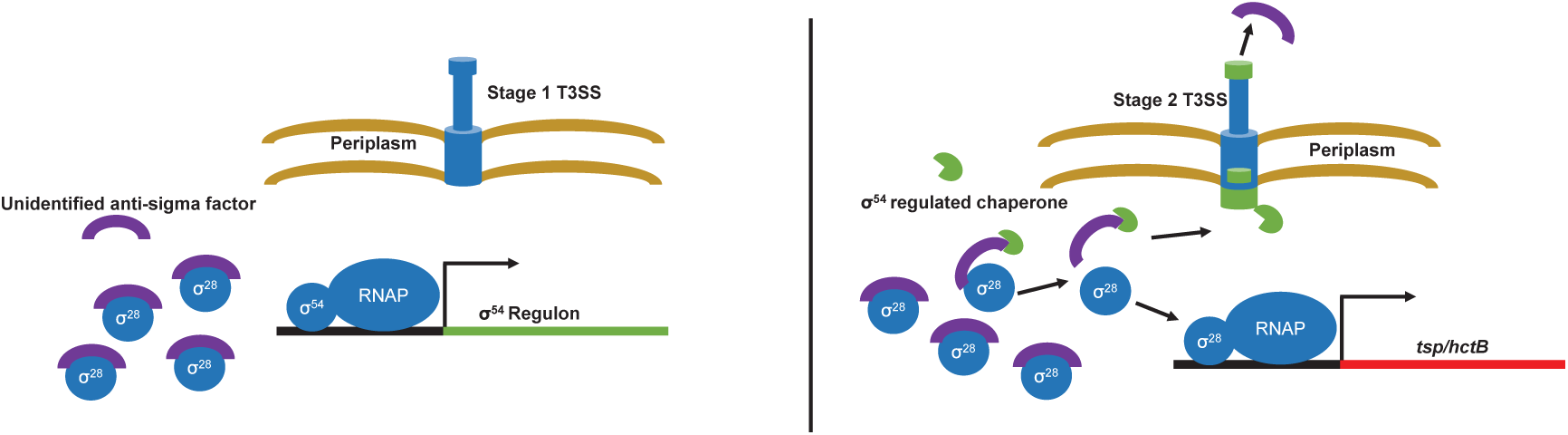
Schematic representation of a hypothetical σ28 regulation cascade. Left) σ28 activity is inhibited during mid-cycle development by an unidentified anti-sigma factor. During the transition to late-cycle, stimulation of the AtoS/C signaling cascade allows σ54 to begin transcribing genes within its regulon. Right) Direct and indirect products of σ54 culminate in the modulation of existing T3SS apparatuses and translation of an unidentified chaperone acting as an anti-anti-sigma factor, thereby releasing σ28 and guiding the anti-sigma factor to be secreted. The liberated σ28 is now free to associate with RNAP and begin transcribing genes within its regulon.

It is important to acknowledge the limitations of our study. Most importantly, RNA-seq in combination with targeted sigma factor dysregulation does not allow distinction between direct and indirect effects. Although combining both overexpression and knockdown approaches narrows this gap, to truly make this distinction, a more direct, chromatin-immunoprecipitation (ChIP) and sequencing-like approach is necessary. Due to the lack of commercially available ChIP-grade chlamydial antibodies, the most reliable methodology would require endogenous tagging of sigma factors using allelic exchange. While allelic exchange has been successfully adapted for Ctr, we have been so far unable to generate a strain with an epitope-tagged σ54. Additionally, expression of the dCas12 protein itself is metabolically taxing, as it appears to slow development ~1 division cycle by 24 hpi if induced at 4 hpi. This creates an “apples to oranges” situation where late-cycle genes in general will appear slightly downregulated due to the delay in development when compared to uninduced samples. We attempted to alleviate this concern by comparing targeted knockdown results to their respective uninduced samples as well as to the induced non-targeting sample. This approach minimizes false positives in exchange for an increased vulnerability to false negatives. For example, *ctl0812* is significantly upregulated during σ28 overexpression but does not appear in our final knockdown data set. While transcripts are reduced almost 3-fold during σ28 knockdown compared to uninduced levels, induction of the dCas12 system alone (NT) appears to reduce *ctl0812* transcripts by 1.6-fold. In turn, our comparison between induced σ28 knockdown and the induced dCas12 alone demonstrated a 1.8-fold decrease and an FDR far above our cutoff (0.54), preventing the gene’s appearance on our final list. Furthermore, several genes have been demonstrated to be regulated by more than one promoter (14, 41, 42). In the case of sigma factor knockdown, it is possible that some genes are transcribed enough from a secondary promoter to be overlooked during statistical analysis. This may also explain discrepancies between our σ54 knockdown data and the Soules et al. constitutively active strain (14), where tandem promoters could explain conflicting data. Again, a ChIP or ChIP-like approach would be ideal for reliably determining the direct regulation of any sigma factor, and we stress our position that our data cannot distinguish between direct and indirect effects. For example, *scc2* (*lcrH_1*) has been previously shown to be σ66 regulated, despite being one of the most significant hits when knocking down σ54 (43).

Yu et al. performed a promoter-prediction analysis alongside *in vitro* validation and determined 10 genes were likely directly regulated by σ28 (13). Our *in vivo* overexpression model was able to recapitulate three of those (*yebL, hctB,* and *tsp*) while our knockdown model only recapitulated *hctB* and *tsp.* Why *yebL* did not appear to be downregulated upon σ28 knockdown is unclear. One plausible explanation, as discussed previously, is the potential for tandem promoters – one recognized by σ28 and a second recognized by σ66. In the event of σ28 downregulation, there may be sufficient σ66 facilitated transcription to prevent a statistically significant decrease in *yebL*. Another possibility is that *yebL*’s increase in transcription was coincidental – given the large disparity between the magnitudes of overexpression, this is not an unreasonable proposition. It is not surprising that the other genes proposed by Yu et al. were not detected – promoter prediction analyses and *in vitro* transcription experiments lack the complex environment in which sigma factors exist, potentially permitting unnatural interactions - i.e., false positives. *Chlamydia* is renowned for the extent to which it has taken reductive evolution; however, this makes it peculiar to claim σ28 regulates only two genes. Why would an organism with such a highly reduced genome maintain a sigma factor that is on the verge of being redundant? Notably, the environmental *Chlamydiae* lack σ28 altogether, making it reasonable to speculate that the σ28 regulon in pathogenic species is as specific as it appears while simultaneously making the retention of it even more puzzling (44). A small regulon is more permissive to sigma factor loss than a large regulon, requiring only a few mutations to permit σ66-dependent regulation and render σ28 non-essential. It is unclear what specific advantage is conferred by having such precise control over HctB and Tsp levels that is not relevant for the environmental *Chlamydiae.* That being said, we have observed that *Chlamydia* is extraordinarily sensitive to alterations in Tsp levels (45).

Similar to the Yu et al. study (13), Mathews and Timms utilized a promoter prediction program to attempt the characterization of the σ54 regulon (12). While we were unable to recapitulate any of those predicted genes, Soules et al. did find two predicted genes in their data set (14). The discrepancy between our study and Soules et al. could be due to the potential presence of tandem promoters, or due to the model design in general. Soules et al. relied on the σ54 EBP, AtoC, lacking a regulatory domain. This mutated EBP was constitutively active, constantly hydrolyzing ATP and hyperactivating σ54. A significant decrease in available ATP can skew the Rsb/σ66 regulatory pathway, creating a potentially confounding environment (29). These deficiencies reflect the reality of the constraints associated with working with obligate intracellular pathogens and the molecular difficulties associated with the scientific question at hand. Furthermore, it provided valuable information to the field, despite the limitations mentioned. Together, our investigations provide *in vivo* experimentation of both the overactivity and knockdown of σ54 in Ctr, providing the field with the most rigorous and comprehensive interrogation of σ54’s potential regulon to date. While both of our studies provide strong evidence for the expanded (direct/indirect) regulon of σ54, more specific experimental techniques are needed to truly define the direct regulon of σ54.

In this first published use of multiplex CRISPRi mediated knockdown in Ctr, we sought to knockdown both alternative sigma factors simultaneously to interrogate any potential cooperative effects. Notably, double knockdown resulted in more significant hits than both individual knockdowns combined. It is unclear why an expansion in significant hits is observed, considering the apparent sequential action of σ54 followed by σ28; one would presume that σ54 knockdown alone would phenocopy a double knockdown. One possibility is that the alternative sigma factors exhibit a low level of cooperative function – e.g., initial σ54 activity may allow σ28 to become active, increasing HctB levels and altering chromosomal topology in a way that promotes additional σ54 activity. While the field generally references “late gene transcription” as a homogenous phase, it is likely more complicated than that. It has been demonstrated that an “early-late” phase exists, in which genes such as *hctA* are transcribed before other canonical late genes (46), and it is reasonable to posit a second phase that includes initial σ54 activity – activity focusing on prepping the T3SS apparatus and necessary effectors. If our model is correct (Figure 3), then action of this phase may gradually free σ28, ushering in a third phase in which HctB and Tsp make significant changes to the chromosome and periplasm topology, respectively. During this chromosomal remodeling, the σ54 regulon may shift slightly.

Overall, our data indicate the alternative sigma factors σ28 and σ54 are dispensable for early and mid-cycle development but are both necessary for late-stage development and the production of infectious EBs. Furthermore, RNA-seq data suggest the σ28 regulon is limited to two late genes, *hctB* and *tsp.* Sequencing data also suggest that endogenous σ28 requires σ54 activity in order to become functionally active. Continued investigation of how σ54 indirectly modulates σ28 activity, either by regulating levels of a candidate anti-sigma factor or by another mechanism, is needed to fully understand the molecular underpinnings of transcription during secondary differentiation. Importantly, this study demonstrates successful implementation of multiplex CRISPRi and its utility for genetic dissection of complex and cooperative regulatory pathways in *Chlamydia*.

## Materials and Methods

### Organisms, cell culture, and chemicals

McCoy cells, a mouse fibroblast cell line, were routinely cultivated at 37°C with 5% CO_2_ in Dulbecco’s modified Eagle medium (DMEM; Gibco, Dun Laoghaire, Ireland) supplemented with 10% FBS. *C. trachomatis* serovar L2 - pL2 (plasmid free) strain (a kind gift of I. Clarke) was propagated and harvested from infected McCoy cell cultures at 37°C with 5% CO_2_. Purified EBs were titered for infectivity by determining inclusion-forming units (IFU) on fresh cell monolayers. All bacterial and eukaryotic cell stocks were confirmed to be *Mycoplasma* negative using the LookOut Mycoplasma PCR Detection Kit (Sigma, St. Louis, MO). Molecular biology reagents were purchased from ThermoFisher unless otherwise noted.

### Individual and multiplex crRNA design

For construction of single and double knockdown plasmids, 2ng of the gBlock(s) (IDTDNA; Coralville, IA) listed in Supplemental Table 4 were combined with 25ng of BamHI-digested, alkaline phosphatase-treated pBOMBL12CRia(e.v.)::L2 for use in a HiFi reaction according to the manufacturer’s instructions (New England Biolabs (NEB); Cambridge, MA). For the triple knockdown crRNA insert, 0.25ng of each dual-targeting gBlock for *fliA* and *incA* was mixed with 0.1ng of each complementary oligonucleotide targeting *rpoN* with overlapping basepairs for the dual knockdown gBlocks (primers sigma54_crRNA/(sigma28)/5’ and sigma54_crRNA/(incA)/3’) for use in a HiFi reaction. Subsequently, 2μL of this HiFi reaction was used as a template for a PCR reaction using primers Bam_Flank_Left/(pBOMBL12CRia)/5’ and Bam_Flank_Right/(pBOMBL12CRia)/3’ and the following cycling parameters with Phusion DNA polymerase: 98°C for 2’ followed by 30 cycles of 98°C for 15 seconds, 64.6°C for 30 seconds, and 72°C for 10 seconds. A final incubation step of 30 seconds at 72°C was performed. A second PCR was performed using 2μL of the first reaction and using the same cycling conditions. The product of the correct size (~315bp) was gel-purified and used in a HiFi reaction with the BamHI-digested, alkaline phosphatase-treated pBOMBL12CRia(e.v.)::L2 as described above. All plasmids were transformed into NEB 10-beta cells and verified by both restriction enzyme digestion and Sanger sequencing prior to transformation into *Chlamydia trachomatis*.

### Transformation of *C. trachomatis*

Transformations were performed using a protocol described previously, with modifications (31). Briefly, for 1 well of a 6-well plate, 10^6^ *C. trachomatis* L2 - pL2 EBs were incubated at room temperature in 50 µL of Tris-CaCl_2_ with 2 µg of plasmid. McCoy cells seeded the prior day were infected using the transformation solution and 2 mLs of Hank’s Balanced Salt Solution (HBSS; Gibco). Cells were centrifuged at 400 x g for 15 minutes at room temperature. Cells were then incubated at 37°C for 15 minutes before HBSS was aspirated and replaced with DMEM. At 8 hpi, 1 µg/mL of cycloheximide and 2 U/mL of penicillin were added to the culture media. The infection was passaged every 40-48 hours until a penicillin resistant, GFP positive population was established.

### Inclusion Forming Unit Assay

McCoy cells were infected at an MOI of 0.3 with relevant strains and induced or not with 10 nM aTc at 4 (KD) or 14 (OE) hpi and allowed to progress until 24 hpi. At 24 hpi, samples were harvested by scraping cells in 2 sucrose-phosphate (2SP) solution. Samples were lysed via a single freeze-thaw cycle, centrifuged at 17k xg for 30 minutes, resuspended (no liquid change), serially diluted, and used to infect a fresh McCoy cell monolayer. Titers were enumerated by GFP fluorescence the following day.

### Immunofluorescence assay

McCoy cells were cultured on glass coverslips in 24-well tissue cultures plates and infected with relevant strains at an MOI of 0.3. All cells were fixed in 100% methanol. Organisms were stained using a primary goat antibody specific to *C. trachomatis* major outer membrane protein (MOMP) and a donkey anti-goat secondary antibody conjugated to Alexa Fluor 488 (Jackson Labs, Bar Harbor, Maine). Images were acquired on a Zeiss AxioImager.Z2 equipped with an Apotome2 using a 100X lens objective.

### Nucleic acid extraction and enrichment from *Chlamydia*

McCoy cells plated in 6-well plates at a density of 10^6^ per well were infected with *C. trachomatis* serovar L2 transformed strains at an MOI of 0.3 for RT-qPCR analysis or 0.6 for samples destined for RNA seq. Infected cells were induced or not using 10 nM anhydrotetracycline (aTc) at 4 hpi in pL12CRia (knockdown, KD) strains or 14 hpi in pBOMBL (overexpression, OE) strains. KD and OE samples were allowed to proceed until 24 or 18 hpi, respectively. At that time, *C. trachomatis* RNA extraction was performed on infected cell monolayers using TRIzol according to the manufacturer’s instructions (Invitrogen/ThermoFisher). Samples were treated with Turbo DNAfree (Ambion/ThermoFisher) according to the manufacturer’s instructions to remove DNA contamination. RNA seq samples were treated using Oligotex mRNA Mini kit (Qiagen, Hilden, Germany) to remove host mRNA contamination with slight amendments to the manufacturer’s instructions – notably, the supernatant was saved for further processing and the Oligotex-bound mRNA was discarded. Samples were then treated using MICROB*Enrich* and MICROB*Express* (Invitrogen/ThermoFisher) according to the manufacturer’s instructions. Samples were aliquoted and stored at −80°C until submitted to the UNMC Genomics Core. Three biological replicates were collected.

### RT-qPCR

cDNA was synthesized from DNA-free RNA using random nonamers (New England BioLabs, Ipswich, MA) and SuperScript III RT (Invitrogen/ThermoFisher) per manufacturer’s instructions. Reaction end products were diluted 10-fold with molecular biology-grade water, aliquoted for later use, and stored at −80°C. Equal volumes of each reaction mixture were used in 25 µL qPCR mixtures with SYBR green master mix (Applied Biosystems) and quantified on a QuantStudio 3 (Applied Biosystems/ThermoFisher) using the standard amplification cycle with a melting curve analysis. Results were compared to a standard curve generated against purified *C. trachomatis* L2 genomic DNA. Amplification of *16S rRNA* was used to normalize respective transcript data. RT-qPCR results were normalized for efficiency with typical results demonstrating r^2^ > 0.995 and efficiencies greater than 90%. Statistical significance was assessed by comparing induced and uninduced samples via ratio paired t-test, * = P value < 0.05.

### Library preparation and RNA sequencing

Initial quality check of starting RNA assessed via Fragment Analyzer (Advanced Analytical) and Nanodrop. The Final libraries were quantified using Qubit DS DNA HS Assay reagents in Qubit Fluro meter (Life Technologies) and the size of the libraries were measured via Fragment Analyzer. Beginning with 400 ng of total RNA from the sample, RNA-seq libraries were prepared using Illumina Ribo-Zero plus Microbiome (Illumina, Inc. San Diego, CA) following the recommended protocol. Resultant libraries from the individual samples were multiplexed and subjected to 100 bp paired read sequencing to generate approximately 60 million pairs of reads per sample using an Illumina Novaseq 6000 in the UNMC Genomics Core facility. DNA library pool was denatured with 0.2N NaOH. The final loading concentration was 300 pM. The original fastq format reads were trimmed by fqtrim tool (https://ccb.jhu.edu/software/fqtrim) to remove adapters, terminal unknown bases (Ns), and low quality 3’ regions (Phred score < 30). The trimmed fastq files were processed by FastQC (47). *Chlamydia trachomatis* 434/Bu bacterial reference genomes and annotation files were downloaded from Ensembl (http://bacteria.ensembl.org/Chlamydia_trachomatis_434_bu_gca_000068585/Info/Index). The trimmed fastq files were mapped to *Chlamydia trachomatis* 434/Bu by CLC Genomics Workbench 12 for RNA-seq analyses.

### RNA Seq Statistical analyses

Each gene’s read counts are modeled by a separate Generalized Linear Model (GLM), assuming that the read counts follow a negative binomial distribution and were normalized based on transcripts per million (TPM). The Wald test was used for statistical analysis of the two-group comparisons. The false discovery rate (FDR) and Bonferroni adjusted p values were also provided to adjust for multiple-testing problem. Fold changes are calculated from the GLM, which corrects for differences in library size between the samples and the effects of confounding factors. Statistically analyzed data can be found in Supplemental Table 5.

## Data availability

The raw and processed RNA sequencing reads in fastq format have been deposited in Gene Expression Omnibus (GEO: www.ncbi.nlm.nih.gov/geo/), Accession number: GSE230645.

## Acknowledgments

The authors would like to acknowledge Drs. E. Rucks (UNMC), I. Clarke (Univ. Southampton), and H. Caldwell (NIH) for reagents, and Drs. E. Rucks, R. Carabeo, and M. Chaussee as well as members of the Chlamydia Research Group at UNMC for helpful comments and suggestions. This work was supported in part by a CAREER award (1810599) from the National Science Foundation (to SPO) and a Ruth L. Kirschstein National Service Research Award (1F31AI164863) from the NIH/NIAID (to NDH).

The University of Nebraska DNA Sequencing Core receives partial support from the National Institute for General Medical Science (NIGMS) INBRE - P20GM103427-19 grant as well as The Fred & Pamela Buffett Cancer Center Support Grant - P30 CA036727. This publication’s contents are the sole responsibility of the authors and do not necessarily represent the official views of the NIH or NIGMS.

## Supplemental Material

**Supplemental Figure S1.** RT-qPCR analysis of *lepA*. Knockdown strains (x-axis) were induced or not with 10 nM aTc at 4 hpi. Samples were collected at 24 hpi and processed as described in the methods. Uninduced ratios of cDNA:16S are set to 1, induced samples’ cDNA:16S ratios are shown as a relative proportion. E.V.: Empty vector, dCas12 only with no crRNA. NT: Non-Targeting. All samples include two biological replicates.

**Supplemental Table S1.** All significantly upregulated hits observed during σ28 overexpression (OE) are listed alongside the fold changes observed during knockdown (KD) or OE of σ28, σ54, or both σ28 and σ54 simultaneously. “Name” includes the annotated name under the accession file used for annotation. Other common names may be given in ( ).

**Supplemental Table S2.** All significantly upregulated hits observed during σ54 overexpression.

**Supplemental Table S3.** Genes significantly downregulated during dual knockdown (KD) of σ28 + σ54 and the accompanying fold changes observed. Also included are fold changes during σ54 knockdown alone and σ28 + σ54 dual overexpression (OE) for those genes. “Name” includes the annotated name under the accession file used for annotation. Other common names may be given in ( ).

S: Soules et al. observed upregulation of these genes in a constitutively active AtoC model, mimicking σ54 hyperactivity.

W: These genes are upregulated during σ28 + σ54 OE but are not observed in σ28 OE.

**Supplemental Table S4.** List of Plasmids, Strains, and Primers used in the study.

**Supplemental Table S5.** Collated, aligned, statistically analyzed RNAseq data for all strains and conditions.

